# Elevated serotonin in mouse spinal dorsal horn is pronociceptive

**DOI:** 10.1101/2023.08.10.552838

**Authors:** Nathan Cramer, Yadong Ji, Maureen Kane, Nageswara Pilli, Luca Posa, Gabrielle Van Patten, Radi Masri, Asaf Keller

## Abstract

Serotonergic neurons in the rostral ventral medulla (RVM) contribute to bidirectional control of pain through modulation of spinal and trigeminal nociceptive networks. Deficits in this pathway are believed to contribute to pathological pain states, but whether changes in serotonergic mechanisms are pro or anti-nociceptive are debated. We used a combination of optogenetics and fiber photometry to examine these mechanisms more closely. We find that optogenetic activation of RVM serotonergic afferents in the spinal cord of naïve mice produces mechanical hypersensitivity and conditioned place aversion. Neuropathic pain, produced by chronic constriction injury of the infraorbital nerve (CCI-ION), evoked a tonic increase in serotonin concentrations within the spinal trigeminal nucleus caudalis (SpVc), measured with liquid chromatography-tandem mass spectroscopy (LC-MS/MS). By contract, CCI-ION had no effect on the phasic serotonin transients in SpVc, evoked by noxious pinch, and measured with fiber photometry of a serotonin sensor. These findings suggest that serotonin release in the spinal cord is pronociceptive and that an increase is sustained serotonin signaling, rather than phasic or event driven increases, potentiate nociception in models of chronic pain.

**Significance Statement:** Serotonergic neurons of the rostral ventral medulla participate in descending pain modulation by regulating spinal and trigeminal nociceptive circuits. Whether changes in serotonergic mechanisms are pro or anti-nociceptive is debated. We show that serotonin release within the spinal trigeminal nucleus is pronociceptive and that enhanced tonic, but not phasic serotonin release may contribute to sensitization in mouse models of chronic pain. These results further clarify the role of serotonin in nociception and suggest that local inhibition of serotonin release or increase of uptake may be a viable therapeutic approach in treating chronic pain.

## Introduction

Growing evidence indicates that chronic pain is related to abnormalities in top-down pain modulatory brain circuits (reviewed by Ossipov et al., 2014; Ren and Dubner, 2002; Heinricher et al., 2009). These descending pathways exert bidirectional control over nociception and an imbalance in this circuitry towards facilitation of postsynaptic targets may promote and maintain chronic pain (Vanegas and Schaible, 2004; You et al., 2010; Ossipov et al., 2014). Therefore, engaging these descending systems to suppress pain signals at early stages of neural processing may be a highly effective strategy for treating pain, especially chronic pain (Bannister et al., 2020). The most completely characterized descending pain modulating circuit is the periaqueductal gray–rostroventral medulla (PAG-RVM) system (Dubner and Ren, 1999; Fields, 2000; Heinricher et al., 2009; Ossipov et al., 2014). The RVM is the final common relay in descending modulation of pain, integrating inputs from PAG and other subcortical and cortical structures to the dorsal horn, as well as the trigeminal nucleus caudalis (SpVc) (España and Berridge, 2006; Heinricher et al., 2009). There is growing evidence that imbalance between facilitatory and suppressive outputs from RVM to dorsal horn neurons contributes to chronic pain states (reviewed by Ossipov et al., 2014; Denk et al., 2014).

RVM contains several classes of cells that express different transmitters (Millan, 2002; Cai et al., 2014; Zhang et al., 2015). Extensively studied are the ON and OFF cells that respond to acute noxious stimuli with increased or decreased firing rates, respectively (Fields, 2000; Heinricher et al., 2009). A significant proportion of RVM neurons—which are thought to be neither ON nor OFF cells (but see Gau et al., 2013)—express serotonin (5HT), which they release upon dorsal horn neurons. However, very few studies report data on unambiguously identified 5HT-RVM neurons, and insufficient data exist upon which to support a scientific premise regarding the role of 5HT in chronic pain. There exist diametrically opposed views on this critical question supported by conflicting evidence, with some suggesting that 5HT is pro-nociceptive, and others that it is anti-nociceptive (Suzuki and Dickenson, 2005; Wei et al., 2008; Viguier et al., 2013; Ossipov et al., 2014; Bannister and Dickenson, 2017; Cooper et al., 2022; Ganley et al., 2023). For example, we showed that—in animals with pain evoked by chronic constriction injury of the infraorbital nerve (CCI) (Bennett and Xie, 1988; Vos et al., 1994)—suppressing 5HT synthesis in RVM significantly ameliorates pain behaviors and reverses the hyperexcitability of SpVc neurons (Okubo et al., 2013). It is possible that 5HT has different effects on spinal versus forebrain structures involved in pain behaviors (Andersen and Dafny, 1983; Santello and Nevian, 2015). Determining whether 5HT is pro or anti-nociceptive is key to developing evidence-based therapeutics for chronic pain, as existing 5HT modulators have had inconsistent clinical efficacies, and have resulted in significant side effects (Finnerup et al., 2010; Bardin, 2011; Tuveson et al., 2011; Patel and Dickenson, 2016).

Barriers to understanding the role of 5HT in chronic pain are due also to the existence of several types of serotonin receptors; some of these receptors evoke neuronal excitation whereas others produce inhibition; some are presynaptic and others preferentially postsynaptic (Raymond et al., 2001). Importantly, the affinity of these receptors to 5HT varies over orders of magnitude, such that different classes are activated preferentially in response to different [5HT] (Lopez-Garcia, 2006; Viguier et al., 2013). As a result, the effects of 5HT on nociception depend critically on the levels of 5HT. Unfortunately, the levels of tonic or evoked 5HT in the dorsal horn, in either normal or chronic pain conditions, are unknown.

To address this knowledge gap, we compared 5HT dynamics in SpVc before and after CCI-ION. We find that *tonic* levels of 5HT are higher in CCI-ION animals. By contrast, *evoked* 5HT levels are indistinguishable in animals with or without CCI-ION. In uninjured mice, optogenetic release of 5HT from RVM terminals in SpVc evokes tactile sensitivity and conditioned place aversion, consistent with a pro-nociceptive role for 5HT.

## Materials and Methods

### Animals

All animal procedures were reviewed and approved by the Authors’ Institutional Animal Care and Use Committee and adhered to the National Institutes of Health guide for the care and use of laboratory animals and ARRIVE guidelines. We use adult male and female transgenic mice in which Cre recombinase expression is controlled by the *Pet-1* promotor (Scott et al., 2005). Experimental mice were bred in-house from breeding pairs purchased from the Jackson Laboratory (JAX stock # 012712; B6.Cg-Tg(Fev-cre)1Esd/J).

### Viral construct injection

All viral vector injections were performed under aseptic conditions, in a stereotaxic device under isoflurane anesthesia with Rimadyl for postoperative analgesia. We targeted the rostral ventral medulla (RVM) via a craniotomy (∼1 to 2 mm) at 6 mm caudal to bregma, on the midline, and injected 500 nL of AAV5-Ef1a-DIO-ChR2(E123T)-EYFP (Addgene plasmid #35507) or AAV5-Ef1a-DIO-eGFP at a depth of 4.6 mm. The viral vector for the serotonin sensor, AAV9-CAG-iSeroSnFR-Nlgn, was produced by the University of Maryland School of Medicine’s Viral Vector Core (Baltimore, Maryland) using Addgene plasmid #128485. Sensor injections into the spinal trigeminal nucleus caudalis (SpVc) were made by reflecting the musculature over the foramen magnum to expose SpVc at the level of the obex. We injected 500 nL of virus at a depth of 0.5 mm bilaterally.

### Quantification of serotonin levels in brain samples

We used Pet-Cre mice of both sexes, 11 of which had constriction injury of the infraorbital nerve (CCI-ION; see below), and 9 receiving sham surgery. Three weeks after surgery, they were deeply anesthetized, and the brains removed. The spinal trigeminal nucleus caudalis (SpVc) ipsilateral to the CCI-ION was microdissected and rapidly frozen. We combined SpVc of 2 mice (of the same sex and treatment group) for each sample. Samples were kept at -80 °C until assay.

A liquid chromatography-tandem mass spectrometric (LC-MS/MS) method was used for the quantification of serotonin levels in brain samples. Serotonin-d4 was used as an internal standard (IS). Analysis was performed on a Thermo TQS Altis Tandem Quadrupole Mass Spectrometer (Thermo Fisher Scientific Corporation, Waltham, MA, USA) using electrospray ionization (ESI) operated in a positive ion mode, with mass transitions of *m/z* of 177.1 → 160.1 for serotonin and *m/z* 181.1 → 164.1 for the IS(Serotonin-d4). Serotonin concentrations were quantified in the concentration range of 5-5000 ng/g of in brain. Chromatographic separation was achieved on a BEH C18 column (2.1 x 50 mm, 1.7 μm; Waters Corporation, Milford, MA) using 0.1% formic acid in water (A) and 0.1% formic acid in acetonitrile (B) as mobile phase using a 4.2 min linear gradient program at a flow rate of 0.3 mL/min. The gradient program starts at 5% B for 0.75 min, increases to 95% B in 1.5 min; and further increased to 99% B in 2.8 min; the composition was brought back to initial (5%) in 3.2 min and maintained until 4.2 min for re-equilibration. The retention time of serotonin and the IS was 0.50 min. Protein precipitation with acetonitrile was used for the extraction of serotonin and the IS from the samples. 20 μL of brain homogenate (500mg/ml homogenized in ice cold water with 0.1% formic acid) was mixed with 10μL of IS solution (500 ng/mL) followed by addition of 800 μL acetonitrile. After vortex mixing for 2 min and centrifugation for 10 min at 15 000 rpm 4 °C, a 700 μL aliquot of supernatant was transferred into 2ml tubes and dried under a steady stream of nitrogen. The samples were re-suspended in 100 μL of a mixture of mobile phase A and B (1:1 v/v) and 2μL was injected into the LC-MS/MS system. Data collection and analysis were performed using Xcalibur V 2.1 (Thermo Scientific, San Jose, CA). Statistical analysis was performed using a two-tailed t-test with an alpha of 0.05.

### Chronic pain models

#### Chronic constriction of the infraorbital nerve

Male and female mice were anesthetized using a mixture of ketamine and xylazine (i.p) and placed in a supine position on a temperature-controlled heating pad. Using aseptic surgical techniques, the infraorbital branch of the trigeminal nerve was exposed through an intraoral incision and freed from surrounding connective tissue. Approximately 1 to 2 mm from its point of exit at the infraorbital foramen, the nerve was loosely ligated with sterile, 4-0 silk sutures. The incision was closed with VetBond tissue adhesive and the mice were monitored continuously until fully recovered from anesthesia. Daily post-surgical monitoring continued for 5 to 7 days while the mice continued to recover in their home cage.

#### Complete Freund’s Adjuvant (CFA)

Male and female mice previously injected with bilateral injections of AAV9-CAG-iSeroSnFR-Nlgn in SpVc mice were anesthetized with isoflurane and 14 μL of CFA was injected subcutaneously into the vibrissae pads. Injections were made bilaterally using a Hamilton syringe with a 30-gauge needle. We obtained fiber photometry recordings from these mice 3 to 5 days after CFA injection.

### Optogenetic stimulation of 5HT afferents

Five female and 6 male Pet-Cre+ mice received RVM injections of either AAV5-Ef1a-DIO-ChR2(E123T)-EYFP (2 female, 4 male) or AAV5-Ef1a-DIO-eGFP (3 female and 2 male), as described above. These mice were implanted with a 2.5 mm ceramic ferrule, 400um Core, 0.39NA fiber optic probe (RWD Life Science) fixed immediately over the dorsal surface of the cervical spinal cord and secured to the adjacent vertebrae with dental cement. This location was chosen to ensure stability of the probes, while enabling stimulation of ChR2-expressing 5HT afferents that project to caudal segments of the spinal cord.

#### Conditioned Place Aversion (CPA)

Mice expressing either ChR2 or eGFP in serotonergic RVM neurons were placed in a two-chamber apparatus and the time spent exploring each side was recorded for 30 minutes. Animals then underwent 3 days of conditioning during which they would receive 30 mins of optical stimulation in the morning and 30 minutes of sham stimulation in the afternoon in their preferred chamber. Stimulus trains of 10 pulses, 4 ms in duration, at 20 Hz were applied every 10 seconds during stimulation. The mice were connected to the fiber optic system with the light disengaged during sham stimulation. The next day chamber preference was recorded once more for 30 mins and time spent in the paired and unpaired sides was compared with a paired t-test.

#### Mechanical withdrawal thresholds

We measured baseline mechanical withdrawal thresholds over 5 consecutive days 3 weeks after the surgery. To assess tactile sensitivity, mice place on a raised platform with a wire mesh floor and allowed to acclimate to the testing environment for 1 hour prior to testing. We tested the left and right hind paw using the up-down method to determine withdrawal thresholds (Dixon, 1965; Chaplan et al., 1994; Akintola et al., 2017).

During these baseline tests, the mice were connected to the fiber optic stimulation system but without the LED engaged. On day 3, we reassessed withdrawal thresholds 30 minutes after 4 repeated optical stimulation trains: 10 pulses (4 ms duration) at 20 Hz with 10 seconds between trains. Before each testing session, the LED power was calibrated to 5 mW. The test session was repeated 48 hours later to test for reproducibility. We used a one-way repeated measures ANOVA with post-hoc Tukey’s multiple comparisons test for changes in threshold within each group across recording sessions.

### Fiber photometry

We used 5 male and 5 female C57Bl6 mice. Mice were anesthetized with an intraperitoneal injection of 2mg/kg urethane, placed on a stereotaxic frame and a fiber optic probe (400 μM diameter, 0.39NA; RWD Life Sciences) was placed superficially over the right or left SpVc. The fiber optic probe was connected to a RZX10 LUX fiber photometry processor running Synapse software (Tucker-Davis Technologies) through a Doric mini cube (Doric Lenses). LEDs at 465 nm (30 μW) and 405 nm (10 μW) were used for iSeroSnFR excitation and isosbestic control respectively. LED power was verified and calibrated as needed using a digital optical power meter (Thor Labs). Responses to noxious pinch were recorded either naïve (3 female and 2 male mice) or 3 to 5 days after bilateral CFA injection (3 female and 2 male mice). We analyzed the data using customized Python scripts adapted from Tucker-Davis Technologies templates which calculated relative changes in fluorescence. Changes in sensor fluorescence were calculated by subtracting the scaled isosbestic signal from the sensor fluorescence. Event related changes in sensor fluorescence were converted to ΔF/F using the 5 second window prior to each stimulation as baseline. The area under the curve (AUC) for the average response was calculated for each mouse using the AUC analysis function in GraphPad Prism.

We used three mice for *in vitro* fiber photometry verification of iSeroSnFr responsiveness to serotonin. Mice were anesthetized with ketamine/xylazine and 300 μM thick coronal slices through the region of sensor expression were prepared following the method described by Ting *et al* (Ting et al., 2014). For recordings, we placed slices in a submersion chamber and continually perfused (2 ml/min) with ACSF containing (in mM): 119 NaCl, 2.5 KCl, 1.2 NaH2PO4, 2.4 NaHCO3, 12.5 glucose, 2 MgSO4·7H2O, and 2 CaCl2·2H2O. Changes in sensor fluorescence in response to bath application of serotonin or norepinephrine were recorded as described above.

### Statistics and Rigor

Personnel performing behavioral analysis were blind to the experimental conditions. All statistical comparisons were performed using GraphPad Prism software. We used an unpaired t-test for comparisons between two separate groups or a paired t-test for within group comparisons. For analysis of mechanical thresholds that occurred across multiple sessions, we used a 1-way repeated measures ANOVA with Tukey’s test for post-hoc comparisons. In all tests, a p – value less than 0.05 was considered significant. Data in all graphs are shown as mean with 95% confidence intervals unless explicitly stated otherwise in the figure legends.

## Results

### Serotonin release in SpVc is pronociceptive and aversive

Serotonergic neurons in the rostral ventral medulla (RVM) are the primary source for serotonergic afferents in the spinal dorsal horn (Kwiat and Basbaum, 1992). We investigated the contribution of these descending serotonergic projections using optogenetics in awake mice. We implanted a fiber optic probe over the cervical spinal cord in PET-1/FEV Cre mice previously injected with AAV5-ChR-EYFP-DIO to selectively activate RVM serotonergic axons in the spinal cord with trains of optical stimuli. Expression was verified in each mouse by histological examination of GFP expression in RVM (Fig 1A). We also verified that optical stimulation of these neurons *in vitro* at frequencies characteristic of the activity of these neurons (Aghajanian and Vandermaelen, 1982; Allers and Sharp, 2003) (10 pulses at 20 Hz) reliably entrained action potentials (Fig. 1B).

**Figure 1:**
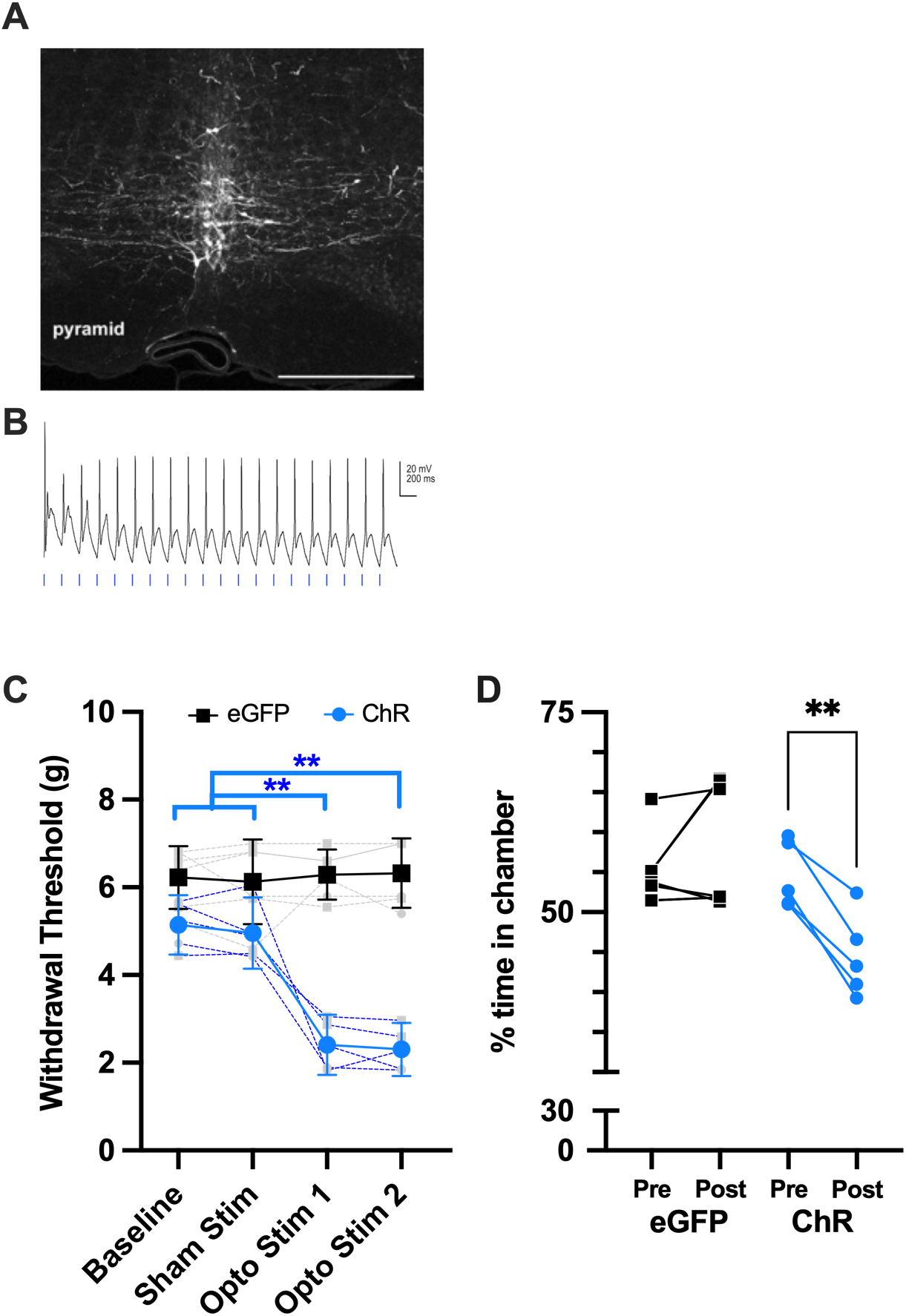
Activation of serotonergic afferents in spinal cord is aversive. (A) Example of serotonergic neurons in RVM transfected to express channelrhodopsin and the fluorescent reporter eYFP. (B) *In vitro* verification that trains of optical stimuli (20 Hz) can entrain ChR expressing neurons in RVM. (C) Optical stimulation (4 repetitions of 10 pulses, 4 ms durations, at 20 Hz with 10 seconds between trains) of spinal serotonergic afferents that express channelrhodpsin produces mechanical allodynia in naïve mice (blue) but not in animals expressing eGFP (black). (D) Optical stimulation of spinal 5HT axons evokes conditioned place avoidance only in mice that express ChR and not eGFP controls. ** = p < 0.01, repeated measures ANOVA in (C) and paired t-test in (D). N = 5 per group.

We assessed the behavioral impact of activating RVM5HT axons on hindpaw nociceptive thresholds using von Frey filaments. Baseline mechanical withdrawal thresholds were calculated for mice that expressed either channelrhodopsin (ChR) or eGFP in serotonergic neurons in RVM (n = 5 per group). Optical stimulation of serotonergic spinal afferents had no effect on mechanical thresholds in the control (eGFP) group but produced mechanical allodynia in mice expressing ChR (Fig. 1C, one-way repeated measures ANOVA with Tukey’s multiple comparison post hoc test, p < 0.01 for baseline vs the first or second optical stimulation). Thus, activation of RVM5HT terminals in SpVc in otherwise naïve mice reliably produces in mechanical allodynia (Fig. 1C).

To examine a non-reflexive behavioral metric, we performed a conditioned place aversion test in the same group of mice. As shown in Figure 1D, mice expressing ChR in RVM5HT axons spent less time in their previously preferred chamber after pairing with optical stimulation, while we observed no change in eGFP controls. As a percentage of the session duration, time spent in the preferred chamber decreased in ChR expressing group from an average of 55% (49 to 60 %) prior to stimulation to 45 % (38 to 51 %; two-tailed paired t-test, p = 0.0019, n = 5) during the stimulation session. As a group, the average time in the preferred chamber decreased by 10 % (6 to 14%) suggesting that activation of spinal RVM5HT afferents in otherwise naïve mice is aversive.

### Serotonin dynamics in SpVc

The findings above suggest that serotonin release in spinal cord is aversive and results in mechanical allodynia. Therefore, we hypothesized that serotonin release would be elevated in mice with injury induced allodynia. We tested this hypothesis using fiber photometry and viral-mediated expression of a genetically encoded serotonin sensor (AAV9-CAG-iSeroSnFR) (Unger et al., 2020) in SpVc. *In vitro*, bath application of 50 to 100 μM serotonin produced dose dependent increases in sensor fluorescence (n = 3 slices) as reported by Unger et al. (example recording in Fig. 2A inset). In contrast, bath application of norepinephrine (20 to 100 μM) did not evoke responses (data not shown).

**Figure 2:**
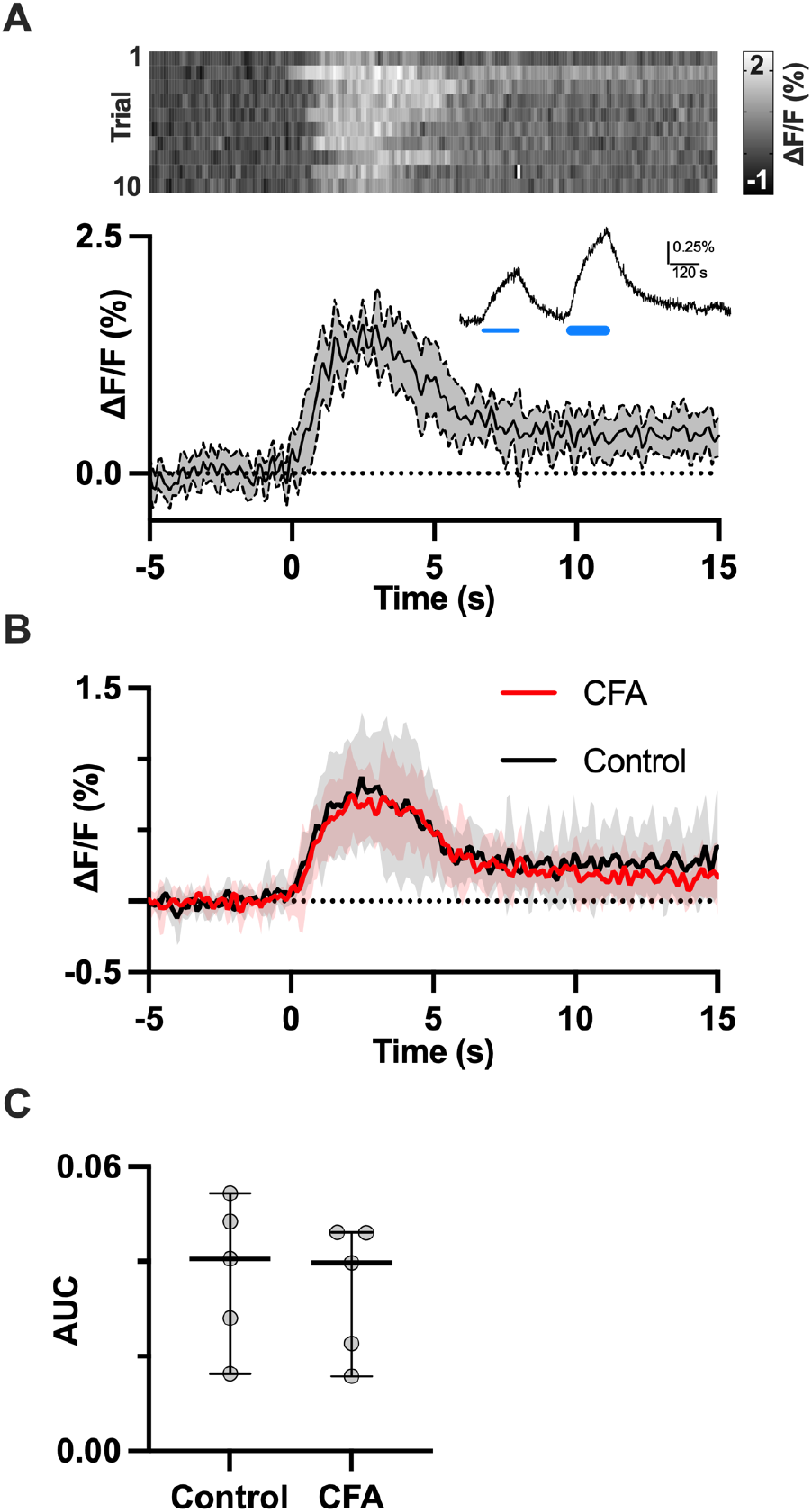
Phasic release of serotonin in SpVc is unaffected by CFA-induced inflammatory pain. (A) Changes in fluorescence of a serotonin sensor expressed in SpVc and measured by fiber photometry in response to noxious heat. The heatmap shows the sensor response to 10 consecutive pinches with the average and 95% confidence intervals of all 10 responses depicted in the trace below. In vitro verification of the sensor response to exogenous application of 50 and 100 μM serotonin (thin and thick blue lines) is shown in the inset. (B) Average of pinch (t = 0) evoked serotonin transients in SpVc from naïve mice and from mice 3 to 5 days after CFA injection (n = 5 per group). There is no difference in the response, measured as area under the curve (AUC), between sham and CFA mice. p > 0.05, t-test.

*In vivo*, fiber photometry recordings from SpVc in urethane anesthetized mice, that were either naïve or had received bilateral subcutaneous injections of CFA into the vibrissae pads, revealed robust serotonin transients in response to noxious pinch. An example of these transients recorded from a sham mouse is shown in Figure 2A, where the heat map shows the responses to individual trials. The line graph below the heat map shows the average response across all trials with 95% CIs indicated by the shaded region. Serotonin levels increased steadily and consistently in SpVc for the duration of the pinch (∼2 s) before returning close to pre-stimulus levels.

Group data from 5 sham and 5 CFA mice (3 female and 2 male mice per group, Figs. 2B and C) revealed no differences in the profile of serotonin release (Fig. 2B) or area under the curve (AUC) between sham and CFA-injected mice (mean AUC with 95% CIs: sham 0.038, 0.018 to 0.057; CFA: 0.034, 0.17 to 0.51, Paired t-test, p = 0.7). The similarity in serotonin transients in SpVc suggests that mechanical allodynia resulting from CFA does not result from differences in serotonin transients triggered by noxious events.

### Tonic serotonin levels

An alternative hypothesis to injury-induced hyperalgesia resulting from larger serotonin transients is that *tonic* levels of serotonin are elevated in mice with chronic pain. We tested this hypothesis using quantitative liquid chromatography-tandem mass spectrometry (LC-MS/MS) to compare tissue concentrations of serotonin in SpVc of mice with a chronic constriction injury of the infraorbital nerve (CCI-ION) and shams controls. Consistent with our hypothesis, concentrations of serotonin in SpVc from CCI-ION mice were higher compared to those in sham mice (Fig. 3; mean ±95% CIs; sham: 0.19 ng/ml, 0.06 to 0.57 ng/mL versus CCI-ION: 0.9 ng/ml (95% CIs: 0.48 to 0.96 ng/mL; unpaired t-test, p = 0.018). Thus, a sustained increase in tonic serotonin levels is associated with CCI-ION injury.

**Figure 3:**
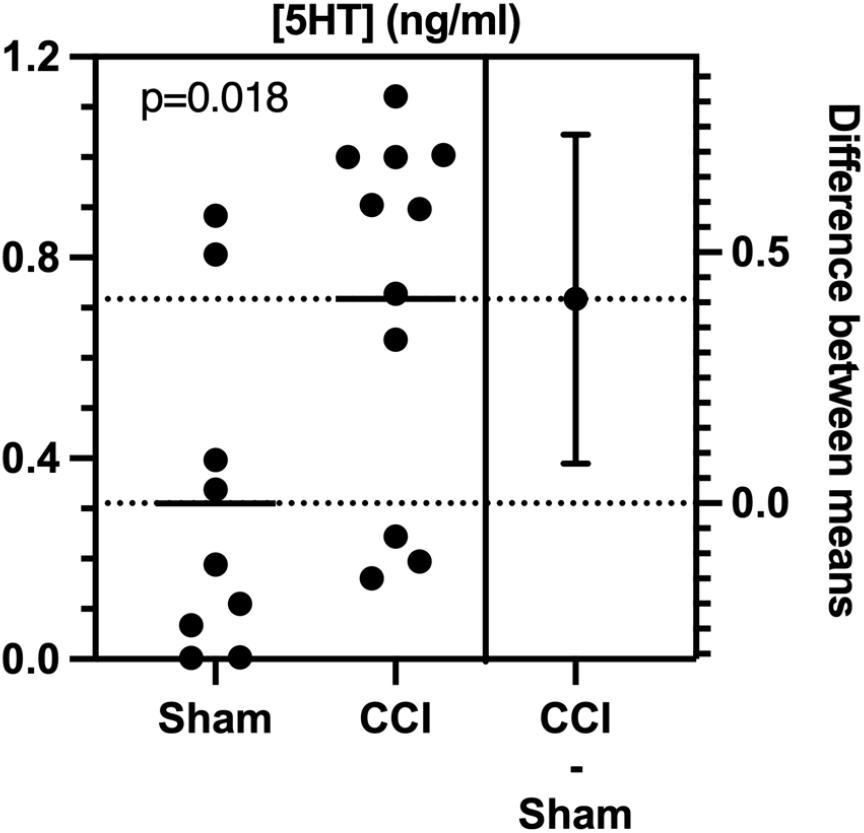
Tonic serotonin levels are elevated in SpVc in a mouse model of neuropathic pain. Concentrations of serotonin in isolated SpVc tissue were quantified by liquid chromatography-tandem mass spectrometry (LC-MS/MS). Group sizes: n = 9 sham and n=11 CCI, p = 0.018, unpaired t-test.

### Sex as a biological factor

Male and female mice were included in each experiment and, where an effect was observed, both sexes were affected. However, we did not have a sufficient number of each sex to directly test whether sex contributes to the effect size.

## Discussion

We investigated the contribution of inputs from rostral ventral medulla (RVM) serotonergic neurons (RVM5HT) to the SpVc in acute and chronic pain, using mouse models of trigeminal neuropathy and inflammation. We report that transient optogenetic activation of RVM5HT terminals in the medullary dorsal horn (spinal trigeminal nucleus caudalis; SpVc) of naïve mice produces mechanical allodynia and is innately aversive. Serotonin transients in SpVc evoked by noxious pinch were indistinguishable in naïve mice and in those with CFA-induced inflammatory pain. However, tonic levels of serotonin were higher in SpVc of mice with trigeminal neuropathy. Together, our data support the hypothesis that RVM5HT neurons exert pronociceptive effects on circuits in SpVc and suggest that a sustained increase of 5HT within the dorsal horn, as opposed to changes in event-driven release, contribute to chronic pain phenotypes.

### Serotonin and acute pain

The RVM is a well characterized link between higher brain regions that participate in descending modulation of pain and nociceptive circuits in the spinal cord and SpVc. Changes in the strength or efficacy of this pathway are believed to contribute to the onset and maintenance of chronic pain (Vanegas and Schaible, 2004; You et al., 2010; Ossipov et al., 2014). RVM contains a heterogeneous population of neurons, including serotonergic neurons in the raphe magnus nucleus reticularis gigantocellularis-pars alpha and the nucleus paragiganto-cellularis lateralis (Kwiat and Basbaum, 1992). RVM5HT neurons may be pro- or antinociceptive through mechanisms that remain poorly understood, and changes in this signaling pathway may contribute to chronic pain (Suzuki and Dickenson, 2005; Wei et al., 2008; Viguier et al., 2013; Ossipov et al., 2014; Bannister and Dickenson, 2017; Cooper et al., 2022; Ganley et al., 2023).

Manipulations that suppress serotonin release in the spinal cord or elevate serotonin concentrations with SSRIs have little impact on nociceptive thresholds in otherwise naïve animals. Targeted suppression of serotonin synthesis in the RVM with RNAi largely depletes the dorsal horn of serotonin but has no effect on baseline mechanical or thermal withdrawal thresholds (Wei et al., 2010). Increases in tonic serotonin levels produced by systemic or intrathecal administration of the serotonin reuptake inhibitor, fluoxetine, also have no effect on baseline pain thresholds (Nemoto et al., 2022). Although the change in tonic spinal serotonin concentrations produced by SSRIs is unknown, these data suggest that the normal behavioral responses to noxious stimuli are insensitive to this enhanced level of serotonergic signaling. These studies suggest that serotonin normally plays, at most, a minor role in modulating spinal nociceptive circuits in naïve conditions. As we show here, this role can change when RVM5HT afferent activity exceeds a high enough threshold. Optogenetic activation of RVM5HT afferents in SpVc produces mechanical allodynia and is aversive in naïve mice (Fig. 1). Similarly, optogenetic activation of serotonergic neurons within RVM produces prolonged mechanical and thermal hypersensitivity in mice, and the duration of hypersensitivity can extend for weeks following repeated activations (Cai et al., 2014). These results support the hypothesis that RVM5HT neurons contribute to the development of chronic pain (Wei et al., 2010), but the mechanisms through which the activity of these neurons begin to facilitate nociception are unknown.

### Serotonin and persistent pain: phasic release

Noxious stimuli in anesthetized rats induces elevations in 5HT that are particularly prominent in the dorsal horn (Weil-Fugazza et al., 1984). We hypothesized that serotonin release in SpVc would be enhanced in conditions of increased pain or sensitivity. Contrary to our hypothesis, the magnitude of serotonin release evoked by noxious stimuli did not differ between naïve mice and mice with inflammatory pain (Fig. 2). Thus, phasic changes in serotonin levels do not appear to contribute to hypersensitivity arising from inflammatory pain. One possibility is that, in animals with inflammatory pain, tonic serotonin levels have increased to a “ceiling” level that cannot be further increased by phasic, noxious stimuli. It is also possible that the primary source of serotonin release shifts from populations of serotonergic RVM neurons that have recently been shown to be anti-versus pronociceptive (Ganley et al., 2023). Our use of a serotonin sensor to detect release would not be able to detect such a shift or the impact of a change in release sites within SpVc on nociception.

That noxious pinch evokes serotonin transients in SpVc appears to be at odds with evidence indicating that RVM5HT neurons do not respond to acute noxious stimuli. Presumptive RVM5HT neurons have been classified as “neutral” cells, in contrast to other neurons that either increase (“ON cells”) or decrease (“OFF cells) their activity during noxious stimulation (Fields, 2000; Heinricher et al., 2009). The apparent discrepancy between these findings may arise from the different metrics used to assess RVM5HT neuronal activity. Whereas we measured serotonin release in SpVc, previous studies measured firing rate of presumptive RVM5HT neurons. It is possible that small changes in activity levels across the population of RVM5HT neurons result in detectable changes in serotonin levels in SpVc. Furthermore, there is evidence that some serotonergic neurons in RVM do, in fact, respond to noxious stimuli (Gau et al., 2013).

### Serotonin and persistent pain: tonic release

Serotonin may facilitate chronic pain through an increase in tonic release. Consistent with this hypothesis, we find that tonic serotonin levels are higher in mice with a chronic constriction injury of the infraorbital nerve (CCI-ION), relative to shams (Fig. 3). This result is in line with reports that depletion of serotonin in RVM5HT afferents prevents the development of allodynia (Wei et al., 2010). Similarly, in a study of patients with osteoarthritis, concentrations of the serotonin metabolite 5-HIAA in cerebrospinal fluid were positively correlated with reported pain severity and impaired conditioned pain modulation (Bjurström et al., 2022). While higher tonic serotonin levels may explain increased pain after injury, shifts in the function of serotonin receptor subtypes may also play a role in the expression of chronic pain. For example, in animals with CCI-ION, the activity of presynaptic 5HT3 receptors is up-regulated, resulting in increased glutamate release from central primary afferent terminals (Suzuki and Dickenson, 2005; Okubo et al., 2013; Kim et al., 2014). However, it is difficult to relate such changes to serotonin anomalies in chronic pain, because serotonin acts on several presynaptic and postsynaptic receptors (Raymond et al., 2001), whose affinity to serotonin varies over orders of magnitude, such that different classes are activated preferentially in response to different [5HT] (Lopez-Garcia, 2006; Viguier et al., 2013). As a result, the effects of 5HT on nociception depend critically on the levels of 5HT.

Here, we report that, in a chronic pain model (CCI-ION), tonic levels of serotonin in SpVc are increased, in parallel with increased pain behaviors. By contrast, the levels of serotonin evoked by noxious stimuli in animals in with CFA injections are indistinguishable from those in control mice. We also show that optogenetically-evoked release of serotonin from RVM terminals in SpVc evokes pain-like behaviors. Although understanding the final impact of the sustained increase in serotonergic signaling on nociception is complicated by changes in expression patterns of receptor subtypes, our study suggests that targeted therapies that reduce the output of spinally projecting RVM5HT neurons may prove effective in reducing the severity of chronic pain.

## Acknowledgments

Research reported in this publication was supported by the National Institute of Neurological Disorders and Stroke of the National Institutes of Health grants R01NS099245 and R01NS069568. Additional support was provided by the University of Maryland School of Pharmacy Mass Spectrometry Center (SOP1841-IQB2014). The content is solely the responsibility of the authors and does not necessarily represent the official views of the National Institutes of Health. The funding sources had no role in study design; the collection, analysis, and interpretation of data; the writing of the report; or in the decision to submit the article for publication.

## Author Contributions

**Nathan Cramer:** Conceptualization, Investigation, Methodology, Formal Analysis, Writing – Original Draft. **Yadong Ji**: Investigation, Methodology, Formal Analysis. **Maureen Kane:** Conceptualization, Investigation, Methodology, Formal Analysis, **Nageswara Pilli:** Conceptualization, Investigation, Methodology, Formal Analysis, **Alberto Castro**: Conceptualization, Methodology, **Luca Posa**: Conceptualization, Methodology, **Gabrielle Van Patten**: Conceptualization, Methodology. **Radi Masri**: Conceptualization, Investigation, Methodology, Formal Analysis. **Asaf Keller**: Conceptualization, Investigation, Methodology, Formal Analysis, Writing – Original Draft, Resources.

